# How to name and classify your phage: an informal guide

**DOI:** 10.1101/111526

**Authors:** Evelien M. Adriaenssens, J. Rodney Brister

## Abstract

With this informal guide, we try to assist both new and experienced phage researchers through two important stages that follow phage discovery, i.e. naming and classification. Providing an appropriate name for a bacteriophage is not as trivial as it sounds and the effects might be long-lasting in databases and in official taxon names. Phage classification is the responsibility of the Bacterial and Archaeal Viruses Subcommittee (BAVS) of the International Committee on the Taxonomy of Viruses (ICTV). While the BAVS aims at providing a holistic approach to phage taxonomy, for individual researchers who have isolated and sequenced a new phage, this can be a little overwhelming. We are now providing these researchers with an informal guide to phage naming and classification, taking a “bottom-up” approach from the phage isolate level.

## 1. Introduction

Virus taxonomy is currently the responsibility of the International Committee on the Taxonomy of Viruses (ICTV, [1]), which published its first report in 1971. The Bacterial and Archaeal Viruses Subcommittee (BAVS) within ICTV holds the responsibility of classifying new prokaryotic viruses. New proposals can be submitted year round by the public. These Taxonomy Proposals (TaxoProps) are evaluated by relevant Study Groups (SGs) and the BAVS [2], and are then discussed and voted on by the Executive Committee (EC) during the yearly meeting. All ICTV-accepted proposals are finally ratified by the members of the IUMS (International Union of Microbiological Societies) Virology Division through an email vote.

Bacterial virus taxonomy has undergone a number of changes since the discovery of bacteriophages in the early 20^th^ century. Electron microscopy, which lead to the recognition of different phage morphologies, and nucleic acid content provided the basis for the first classification scheme [3,4]. Ever since, genome composition and morphology have been the major criterion for classification at the family rank, with the current taxonomy comprising 22 families grouping bacterial or archaeal viruses.

For many years, the grouping of prokaryotic viruses in lower rank taxa such as genus and subfamily, happened at a minimal pace. Taking the tailed phage families as an example, the 5^th^ Report of ICTV (1991) described one genus in each of the families *Myoviridae, Podoviridae* and *Siphoviridae* [5]. This increased to 16 genera spread over the three families by the 7^th^ Report [6] and 18 genera by the 8^th^ Report [7]. As nucleotide sequencing techniques improved the number of publically available bacteriophage sequences, researchers started to question the large number of bacteriophage genomes that remained unclassified [8]. These concerns would prove prescient, and in the coming years next generation sequencing methods would spur an explosion in bacteriophage sequencing (Figure 1).

**Figure 1.**
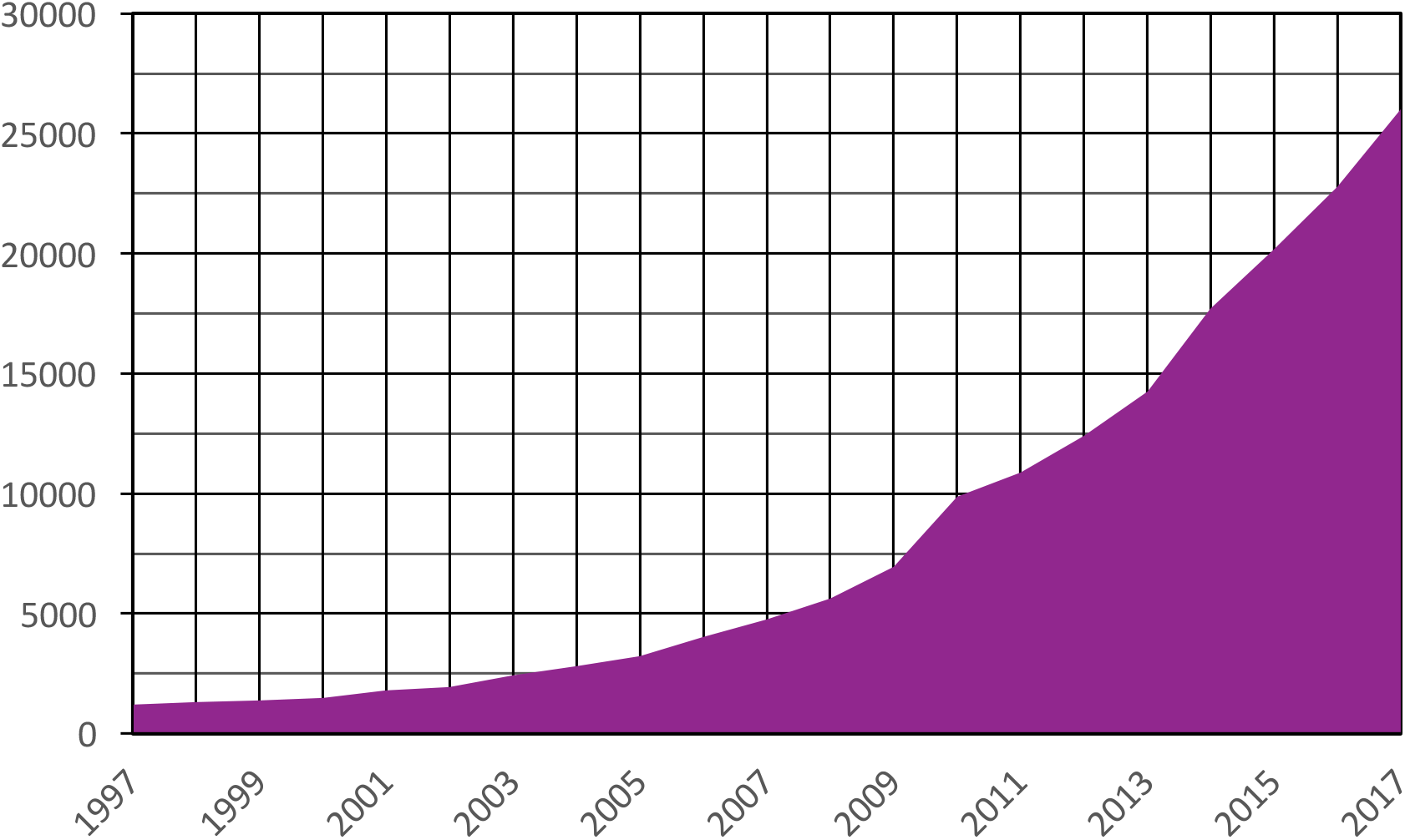
Number of bacteriophage nucleotide sequences deposited in INSDC databases. The number of nucleotide sequences publically available from INSDC databases was calculated by searching the GenBank nucleotide database with the term “vhost bacteria[filter]” and plotting the number of sequences available on January 1 of each year shown [9].

The availability of genome sequence data also gave rise to a range of potential classification or grouping schemes, such as the Phage Proteomic Tree [10], phage network clusters [11], kmer-based grouping [12], signature genes-based grouping [13] or whole genome nucleotide identity-based grouping [14], which were not always compatible with the rules laid out in the ICTV Code and/or the International Code of Virus Classification and Nomenclature (ICVCN). Since the 8^th^ Report of ICTV, both genome and proteome-based methods have been used by the BAVS to classify phages into species, genera and subfamilies, resulting in 14 subfamilies, 204 genera and 873 species in the 2015 taxonomy release [15–20].

In this paper, we provide a naming and classification guide for researchers who have isolated and sequenced a novel bacteriophage isolate specifically, however, these guidelines can be applied to archaeal viruses as well. The guide will follow a “bottom-up” approach, i.e. starting at the species level, rather than the “top-down” approach which was used in the past to assign isolates to a family based on morphology.

## 2. A short, informal guide to naming and classifying your phage

### 2.1. Congratulations, you have isolated a bacteriophage!

You have just isolated and sequenced a bacteriophage, so what do you do next? Well, hopefully, you plan on publishing your finding and submitting the sequence to one of the public databases that are part of the International Nucleotide Sequence Database Consortium (INSDC), GenBank, the European Nucleotide Archive (ENA) or the DNA Data Bank of Japan (DDBJ) [21]. That's great, but now the big question is: “What do I name my phage?” Didn't think about that one, did you? Well, turns out the name you give to your new virus isolate is pretty important, as it will be used in publications, mentioned in conversations among colleagues, and identify the sequence of your virus in public databases and other resources.

Perhaps the most important rule of bacteriophage naming is “don't use an existing name.” There are already four dissimilar bacteriophages named N4, making it very difficult to distinguish between them. So before naming your bacteriophage – and definitely before publishing a report on it – please take the time to compare proposed names against those already used within the field. A good, if not dated, place to start is Bacteriophage Names 2000 [22]. A more up to date list of bacteriophage names can be found by searching the NCBI Nucleotide database [23] with the term “vhost bacteria[filter] AND ddbj_embl_genbank[filter]” [9]. This search will return all bacteriophage isolate names currently associated with sequences in INSDC databases– both those classified by ICTV, as well as those that have yet to undergo official classification.

The current approach to bacteriophage naming is a tripartite construct consisting of the bacterial host genus name, the word “phage,” and a unique identifier, for example “Escherichia phage T4.” Since the first two components of this naming construct are not unique, the third component is critical to the usability of the name. Leafing through a list of bacteriophage names, it is clear that there are a number of approaches to providing unique identifiers in names. For example, one approach to constructing unique identifiers includes information about phage morphology and host [24]. So the name Escherichia phage vB_EcoM-VR20 denotes a **v**irus of **B**acteria, infecting ***E****scherichia* ***co****li*, with myovirus morphology. One caveat to this approach is that one needs to employ electron microscopy or computational methods to derive the correct morphotype. While there are few hard and fast rules for these terms, please be careful when choosing one, because it is likely to be used as shorthand in a variety of contexts for years to come.

Please use the following bacteriophage naming guidelines:

- Always use the complete host genus name, followed by a space, followed by the word “phage,” followed by a space, followed by a unique identifier, e.g. Escherichia phage T4.
- Use only the isolation host genus in the name, rather than higher order taxa names – such as Enterobacteria, Pseudomonad, or the generic Bacteriophage - or lower order taxa names like *Staphylococcus aureus* DSM 1234.
- Do not combine the host genus and the word “phage” into a single word, for example, Mycobacteriophage, Mycophage, etc.
- Do not use an existing unique identifier in the name.
- Do not use Greek letters in the unique identifier.
- Do not start the unique identifier with a numeral and do not use only use only a single letter. Identifiers should include enough complexity to easily distinguish your bacteriophage form others.
- Do not use hyphens, slashes or any type of special character like %$@ etc. You may use underscores to separate parts of the designation, for example vs_p123_233, but these underscores cannot be carried over into official taxon names (see paragraph 2.4).
- Do not use controversial names/phases, profanity, names of prominent people, and trademarked names/phrases as unique identifiers.
- Please do contact the friendly folks on the BAVS if you have any questions.

### 2.2 What is the relationship between bacteriophage isolate names and taxa names?

The rules for naming taxa are described in the International Code of Virus Classification and Nomenclature [25]. Typically, the name of a species is based on the name of the first sequenced isolate, which then becomes the type isolate. Current bacteriophage species names replace “phage” in the tripartite isolate name construct with “virus,” so the isolate Escherichia phage T4 belongs to the species *Escherichia virus T4*. Higher order taxa names are derived from unique identifiers used in isolate names as in the genus *T4virus*. Sometimes these unique identifiers are too similar to existing taxon names, inappropriate, or do not otherwise conform to ICTV taxa naming standards, and a different taxon name must be chosen.

### 2.3 Now it is time to publish your phage sequence

Once you have isolated, sequenced, and named your new bacteriophage, it is time to start thinking about sharing your data with the world. Today, sharing your results is not simply about publishing in a peer reviewed journal. While such descriptions are central to the scientific process, so too is the sequence of your new bacteriophage. Though often overlooked, submitting your new bacteriophage sequence to a public INSDC database such as GenBank, is critical to making your sequence publically available for generations to come. Please keep in mind that in this age of bioinformatics and computational biology, it is likely that over time the sequence record for your new bacteriophage will be accessed exponentially more often than a traditional publication. In other words, do your best to provide detailed and accurate information about your phage when you submit the sequence to an INSDC database. This includes providing the most accurate classification data possible.

If known, lineage information should be included in INSDC sequence submissions using the “lineage” field or in a source note. For example, if you have sequenced a new phage that belongs to the species *Escherichia virus T4*, provide the name of your new virus, e.g. “Escherichia phage My_New_Virus,” and the lineage “Viruses; dsDNA viruses, no RNA stage; Caudovirales; Myoviridae; Tevenvirinae; T4virus; Escherichia virus T4”. In cases where your new phage cannot be placed in an existing species, provide a lineage that reflects classification. For example, if your new phage belongs to the genus *T4virus*, provide the lineage “Viruses; dsDNA viruses, no RNA stage; Caudovirales; Myoviridae; Tevenvirinae; T4virus".

Please use the following guidelines when submitting to public databases:

- Do include lineage information for all submitted sequences. Even if your bacteriophage is novel and does not belong to a described species, provide the most accurate lineage information possible that places the sequence including genus and/or family using the criteria discussed in this manuscript.
- Do include accurate genomic composition information when no other lineage information is available or can be inferred. In most cases it should be possible to place a new isolate within the higher order dsDNA, ssDNA, dsRNA, or ssRNA lineage groupings.
- Do identify prophages using the “proviral” location descriptor.
- If you have questions about sequence submission to INSDC databases, please see The GenBank Submissions Handbook [26].
- If you still have questions, contact GenBank or another INSDC database.

### 2.4 Classifying bacteriophage

So why does taxonomy even matter? Well, taxonomy offers a very useful way of aggregating genome sequence data around a collection of genetic and/or molecular attributes. In this way, rules describing taxa are effectively search terms that allow you to retrieve a set of sequences with similar characteristics. Taxonomy also provides context to sequences when searching for sequence similarities. Knowing that a newly sequenced virus is highly similar to a previously classified one, immediately tells you something about the new virus – the expected gene content, host range, environmental niche, etc.

Bacteriophage classification also supports the organization of genome sequence data within public databases. Each viral species is represented in by at least one “reference” genome in the NCBI Viral Refseq database. Other validated genomes belonging to the same species will be stored as so called “genome neighbors” of the RefSeq genome [9,27]. This arrangement provides a compressed dataset where each species is represented by one (or more) representative sequences – typically from type isolates - as well as a method for retrieving a set of validated genomes from each viral species.

#### 2.4.1 Does my phage represent a new species?

The first question you need to answer is basic one: “Does my newly sequenced phage belong to an existing species?” The main species demarcation criterion for bacterial and archaeal viruses is currently set at a genome sequence identity of 95%, meaning that two viruses belonging to the same species differ from each other by less than 5% at the nucleotide level. This can be calculated by comparing your sequence to existing phage genomes. There are several tools to do this (e.g. BLASTN [28], PASC [29], Gegenees [30] or EMBOSS Stretcher), but each comparison needs to be checked for genomic synteny. While it is common for larger dsDNA phages to differ in their genome organization, isolates showing high levels of rearrangements can never belong to the same species.

If your phage belongs to an existing species, be sure to specify that taxonomic lineage when depositing the sequence into GenBank or other INSDC databases. If results suggest that your phage represents a novel species, congratulations! The appropriate Study Groups and BAVS will, with your help in providing data, create a new species based on your phage. To place this new species in a higher taxon, we will move up to the genus level in the next section.

We recommend that you alert the appropriate BAVS Study Group chair or the Subcommittee Chair [2] who will advise you on how to proceed. This will generally involve filling out an ICTV Taxonomy Proposal Submission Template (TaxoProp for short) which is available here from the ICTV website [31]. The ICTV website includes examples of completed TaxoProps.

#### 2.4.2 Is my phage a member of an existing genus?

The BAVS currently describes a genus as a cohesive group of viruses sharing a high degree of nucleotide sequence similarity (> 50%), which is distinct from viruses of other genera. For each genus, defining characteristics can be determined, such as average genome length, average number of coding sequences, percentage of shared coding sequences, average number of tRNAs, and the presence of certain signature genes in member viruses. The latter can in turn be used for phylogenetic analysis with other bacterial or archaeal viruses encoding this gene to find monophyletic groups as well as higher order relationships.

All the genera currently in the ICTV database have a taxonomy history (TaxoProp) accessible through the website, which can be used for researchers to assess the genus inclusion criteria. If your phage falls into an existing genus, the BAVS will define the new viral species within the existing genus. If the phage is sufficiently different from existing isolates, we can define a new genus, according to the characteristics described above. The minimum requirements for the creation of a new genus are the description of the average genome characteristics of its proposed members (size, GC content, tRNAs, coding sequences), a nucleotide identity comparison with visualization, a comparison of the predicted proteomes and phylogenetic analysis of at least one conserved gene, all of these with the appropriate outliers to demonstrate cohesiveness of the new genus.

While you can propose a new genus and species based upon a unique virus, the BAVS generally recommends that genera be established when two or more related viruses have been deposited in one of the INSDC databases.

#### 2.4.3 What about subfamilies and families?

In the current taxonomy releases, bacterial and archaeal viruses are classified at the family rank according to the morphology of their virions, e.g., phages with short tails are placed in the family *Podoviridae*. This means that for proper classification, electron micrographs of the viral particles should be taken. Based on the morphology and the genomic information necessary for classification in species and genus, we can now look whether your isolate falls in an existing subfamily of viruses. If your new phage, in its newly created genus, is genomically or proteomically similar to phages in an existing subfamily, the genus can be added to the subfamily. The criteria for inclusion can vary between subfamilies and should be consulted from the TaxoProps describing the respective subfamily.

At this time, subfamilies are only created when they add necessary hierarchical information (ICVCN Rule 3.2). In practical terms, this mean that a new subfamily is created when two or more genera show an obvious relation which is not adequately described at the family level. For instance, in the family *Podoviridae*, the subfamily *Autographivirinae* groups all podoviruses that contain an RNA polymerase gene in their genome [16]. The requirements for the creation of a new subfamily are not easily defined, but should definitely include the description of at least two clearly related genera within a family, with evidence that the new subfamily is cohesive.

In a genome-based taxonomy, the tailed phage families *Myoviridae, Siphoviridae* and *Podoviridae*, have become an artificial “ceiling” prohibiting the accurate description and grouping of the genomic diversity present among their member phages. For example, T4-related phages, infecting a wide range of host bacteria from different phyla, are characterized by the presence of a set of 30 conserved (core) proteins [32], but also have more distant cousins in the Far-T4 group sharing only 10 core proteins [33]. These phages are currently all classified within the family *Myoviridae*, but have the genomic diversity to represent a new order. Another example involves the “lambdoid phages”, comprising both siphoviruses (Escherichia phage Lambda) and podoviruses (Salmonella phage P22), which cannot be grouped together in the same family at this time. The BAVS is therefore working on a new system that would abolish these families in favor of genome/proteome-based family descriptions.

#### 2.4.4 My phage/virus does not fit in anywhere, what now?

In the very special circumstance that your new phage does not fit in with any known bacterial or archaeal virus, genomically or morphologically, it is the first representative of a new family. In this case, we strongly urge you to contact the BAVS Chair, or the chair of an appropriate Study Group, to work together to define the demarcation criteria for this new family.

### 2.5 Proposed software to use

This is a non-exhaustive list with suggested software to use. The BAVS as a subcommittee is not associated with the developers of the software described below.

- Nucleotide sequence comparison: NCBI BLASTn [28], Gegenees (uses BLASTn) [30], PASC [29], Gepard dotplot [34].
- Comparison of protein groups, predicted proteomes, identification of signature genes: CoreGenes 3.5 [35,36], Roary (core and accessory genome analysis) [37], prokaryotic Virus Orthologous Groups resource (pVOGs) [38].
- Multiple alignment and/or phylogenetic analyses: Clustal Omega [39], MUSCLE [40], phylogeny.fr [41], MEGA6 [42], FastTree [43].
- Visualization: progressiveMAUVE [44], Easyfig [45], BRIG [46].

## 3. Conclusion

Bacteriophage genomics, ecology, and evolution are quickly growing fields, and large numbers of new phages are being discovered, named, sequenced, and deposited into public databases. This poses semantic, logistical and taxonomical challenges that we have tried to address in this informal guide. It is also important to understand that taxonomy is ever changing because of the unremitting flow of new information. The effort of classification is currently undertaken by a small group of dedicated scientists, but with input from the larger phage community – this means you, dear reader – we can increase the effort while keeping it manageable for each individual researcher.

## Supplementary Material

This article has no supplementary materials associated with it.

## Acknowledgments

We would like to thank Andrew M. Kropinski for his invaluable support and comments on this manuscript, Linda Frisse and Detlef Leipe for discussions about sequence submission guidelines, and Jens H. Kuhn for applying his attention to detail and keen eye to these writings.

## Author Contributions

E.M.A. conceived the study, E.M.A. and J.R.B. wrote the paper.

